# Plasma Micro-RNA Signatures of Type 1 Ryanodine Receptor Related Myopathies

**DOI:** 10.64898/2026.05.14.725164

**Authors:** Pooja Varma, Mayra Saintilus, Morin Nessim, Joshua J. Todd, Payam Mohassel, Tokunbor A. Lawal

## Abstract

Pathogenic *RYR1* variants are associated with a set of rare neuromuscular disorders termed *RYR1*-related disorders (*RYR1*-RD). Clinical manifestations of *RYR1*-RD include proximal/axial muscle weakness, delayed motor milestones, impaired mobility, muscle pain, and fatigue. Muscle-specific microRNAs (miRNAs) are mostly expressed in muscle tissue and can be detected peripherally in plasma. Using a digital detection system, here we identified and quantified differential amounts of miRNAs in six adult (four monoallelic and two biallelic) *RYR1*-RD patient plasma samples compared to controls. Overall, 51 differentially expressed miRNAs were identified and hsa-miR-4454+hsa-miR-7975, in particular, was significantly overexpressed relative to controls (+ 39-fold, P=0.00285). Exploration of these differentially expressed miRNAs warrant further investigation as potential biomarkers of *RYR1*-RD.

## 1. Introduction

The skeletal muscle ryanodine receptor (RyR1) is an intracellular Ca^2+^ release channel that is integral to excitation-contraction coupling (ECC) and is embedded in the sarcoplasmic reticulum membrane.^1 2^ Pathogenic variants in the *RYR1* gene can disrupt intracellular calcium flux and/or cause in decreased expression of RyR1, most often resulting in *RYR1*-related congenital myopathies (*RYR1*-RM) and malignant hyperthermia, an allelic pharmacogenetic disorder.^3^ *RYR1*-RM are estimated to affect approximately 1:90,000 pediatric individuals in the United States, and often have a slowly progressive disease course.^4^ Monoallelic, biallelic and *de novo* cases have been reported.^3^ Clinical manifestations of *RYR1*-RM include hypotonia, exercise intolerance, fatigue, proximal muscle weakness, contractures, scoliosis, ophthalmoplegia, and respiratory insufficiency.^2^ Diagnosis of *RYR1*-RM is often delayed and challenging due to the overlapping disease spectrum, histotype, and the large number of *RYR1* variants of uncertain significance.^5 6^ Currently, clinical management is focused on supportive care as there is no FDA-approved treatment for *RYR1-*RM.

There is expanding interest in microRNAs as potential biomarkers in muscle disorders,^7^ however there is limited data available in *RYR1*-RM.^8^ Identification of novel muscle-related miRNA may reveal common epigenetic therapeutic targets for muscle-related conditions such as sarcopenia, muscular dystrophies and congenital myopathies.^9 10 11^ MicroRNAs (miRNAs) regulate translation using mRNA cleavage or steric hindrance,^12^ intercellular signaling, and are implicated in the pathogenesis of numerous disorders.^13^ Broadly, miRNAs have been linked to disorders of Ca^2+^ dysregulation,^14^ with differential roles in both neurodegeneration and neuroprotection.^15 16^ MiRNAs have also been identified as regulators of RyR expression in neuronal and non-neuronal cell types.^17 18 19 20 21^ Muscle-associated miRNAs are important regulators.^22^ Expression levels of muscle-associated miRNAs predicted to bind *RYR1* 3’ UTR (miR-22, miR-124, miR-128) have been reported.^23 24^

The present study identifies miRNAs in patients with *RYR1*-RM and age and sex-matched healthy controls.

## 2. Subjects and Methods

The exploratory analysis presented here was conducted using deidentified and unlinked baseline (pre-dose) data and biospecimens obtained from individuals enrolled in a prior open-label clinical trial (NCT04141670).^25^ All individuals provided written informed consent under the National Institutes of Health (NIH) Intramural Institutional Review Board (IRB)-approved primary research protocol (20-N-0005) before initiating study procedures and also consented to data and specimens being used for future research. Participants were adults with a diagnosis of *RYR1*-RD. ^25^

### 2.1 Human biospecimen collection and preparation

A total of six *RYR1*-RD affected individuals and six age-matched (±5 years), and sex-matched controls were included in this study. Matched control blood was obtained from healthy donors through the NIH Clinical Center Blood Bank. All samples were obtained by venipuncture, and collected in EDTA tubes. Plasma aliquots were snap frozen on dry ice, and stored at -80°C until analysis.

### 2.2 RNA extraction and miRNA purification/quantification

Prior to extraction, plasma samples were thawed at room temperature and centrifuged at 8,000 xg for ten minutes to separate any debris. Supernatant was removed carefully and transferred to fresh 2 mL Eppendorf DNA LoBind tubes. Isolation of total microRNA from plasma from trial participants (n=6) and healthy controls (n=6) was done using the miRNeasy Serum/Plasma Advanced kit (Cat No./ID: 217204, QIAGEN, Germany) according to the manufacturer’s protocol. RNA was resuspended in 10 μL of RNase-free water and then stored at −80 °C until analysis. RNA concentration was determined using Qubit 3.0 Fluorometer (Life Technologies, USA) and RNA purity was determined using RNA 6000 Pico Kit (Cat No./ID: 5067-1513, Agilent, USA).

### 2.3 Nanostring

The RNA extracted ((3 μL)(n=12)) from plasma samples underwent miRNA expression profiling using the nCounter Human v3 miRNA Expression Assay Kit (Cat No./ID: CSO-MIR3-12, Nanostring, USA). Twelve samples (6 patient and 6 control) were analyzed using a single cartridge. The nCounter miRNA Sample Preparation Kit provided reagents for ligating unique oligonucleotide tags, specific for each of the 800 miRNAs in the panel, onto target miRNAs from the plasma extracted miRNA. Next, ligation, purification and overnight hybridization was followed according to the manufacturers protocol. The miRNAs were hybridized to a capture/reporter probe pair. This capture probe was used for binding the probes to the imaging surface and the reporter probe had a color-coded molecular barcode which was used to identify the specific miRNA targets during detection and scanning.

The Nanostring nCounter system provides platforms for gene expression analysis. nCounter technology is based on a novel method of direct molecular barcoding and digital detection of target molecules using color-coded probe pairs. The nCounter miRNA Expression assays can profile over 800 highly curated human miRNAs (see suppl table 1 – Human_v3_miRNA Gene List) considered biologically relevant for potential clinical application from over 38,000 known sequences in miRBase database (https://www.mirbase.org), and offer direct, digital counts of each miRNA found in a sample.^26^

Using the nCounter SPRINT Profiler, hybridized samples were loaded onto an nCounter Cartridge for processing and data collection. Digital images were captured and processed, and the molecular barcodes found within the samples were counted. Data was extracted to a .csv file to the instrument, then analyzed using nSolver Analysis Software 4.0 and ROSALIND. Counts of more than 50, after background correction, were used to define presence of a given microRNA.

### 2.4 Statistics

The ROSALIND platform was used to identify differentially regulated genes between treatment groups (±1.5 fold-change; p <0.05, standard cutoff for identification of differentially expressed genes). The p-values were adjusted using the Benjamini-Hochberg method. MicroRNAs with a fold change of less than -1.5 or greater than 1.5 were defined as differentially expressed. Log2 normalization was tabulated in order to measure the up/down regulated genes between samples. MicroRNAs with p-values less than 0.05 were considered to have a significant difference in expression between the patient and control samples. ROSALIND was also used to generate a heat map of differentially expressed miRNAs. Blue indicates a lower expression level and orange indicates a higher expression level.

## 3. Results

A total of 6 *RYR1-*RD affected individuals (male n=3, female n=3) had whole blood specimens available for analysis. The patient and control samples were obtained from adults with minimum and maximum ranges between 33 and 49 years old and a median of 39 years old. The *RYR1*-RD patient cohort included biallelic (n=2) and monoallelic (n=4) cases as previously described.^25^

The miRNA analysis, following exclusion of background levels, identified 51 miRNAs to be differentially expressed. Among the 51 miRNAs, 49 were reported to be under-expressed while two, Hsa-mir-4454+hsa-mir-7975 (38.85-fold change, p<0.05), were reported to be over-expressed compared to controls. **Table 1** shows the fold changes and p-values of identified miRNAs in patients over controls. The heat map and volcano plot of the 51 miRNAs identified in patient samples compared to control are depicted in **Figures 1** and **2**, respectively.

**Table 1.** Gene Expression Data.

| Name | Fold Change | p-Value |
| --- | --- | --- |
| hsa-miR-4454+hsa-miR-7975 | 38.8537 | 0.00285 |
| hsa-miR-3168 | -1.52326 | 0.01334 |
| hsa-miR-3161 | -1.54157 | 0.02215 |
| hsa-miR-495-5p | -1.55389 | 0.01758 |
| hsa-miR-3136-5p | -1.62679 | 0.00492 |
| hsa-miR-548y | -1.6963 | 0.0095 |
| hsa-miR-1185-5p | -1.71529 | 0.01054 |
| hsa-miR-627-3p | -1.71638 | 0.01169 |
| hsa-miR-544a | -1.84568 | 0.01463 |
| hsa-miR-411-5p | -1.87945 | 0.01937 |
| hsa-miR-548j-5p | -1.95163 | 0.00193 |
| hsa-miR-105-5p | -1.95262 | 0.003 |
| hsa-miR-548d-3p | -1.96915 | 0.0086 |
| hsa-miR-613 | -1.98052 | 0.00933 |
| hsa-miR-216b-5p | -1.98119 | 0.00815 |
| hsa-miR-4455 | -2.07435 | 0.04029 |
| hsa-miR-561-5p | -2.07865 | 0.03047 |
| hsa-miR-450b-5p | -2.09741 | 0.01475 |
| hsa-miR-548ar-3p | -2.10236 | 0.00188 |
| hsa-miR-644a | -2.14304 | 0.0416 |
| hsa-miR-1253 | -2.18806 | 0.02005 |
| hsa-miR-1285-5p | -2.27327 | 0.00437 |
| hsa-miR-141-3p | -2.27425 | 0.00212 |
| hsa-miR-520d-3p | -2.29842 | 0.02564 |
| hsa-miR-587 | -2.31948 | 0.0016 |
| hsa-miR-32-5p | -2.34088 | 0.00787 |
| hsa-miR-378e | -2.39875 | 0.00082 |
| hsa-miR-206 | -2.48153 | 0.00298 |
| hsa-miR-2116-5p | -2.51924 | 0.01736 |
| hsa-miR-944 | -2.58957 | 0.00097 |
| hsa-miR-590-5p | -2.68395 | 0.0176 |
| hsa-miR-4531 | -2.76157 | 0.00114 |
| hsa-miR-1257 | -2.82555 | 0.00324 |
| hsa-miR-3144-5p | -2.85644 | 0.00178 |
| hsa-miR-873-5p | -2.86056 | 0.0053 |
| hsa-miR-92a-3p | -2.87609 | 0.01439 |
| hsa-miR-941 | -2.91984 | 0.0091 |
| hsa-mir-549a | -2.95634 | 0.00482 |
| hsa-miR-219b-3p | -2.96997 | 0.01115 |
| hsa-miR-548q | -3.01877 | 0.00554 |
| hsa-miR-5010-3p | -3.03888 | 0.00626 |
| hsa-miR-4536-5p | -3.10566 | 0.00634 |
| hsa-miR-1283 | -3.33124 | 0.00072 |
| hsa-miR-656-3p | -3.41506 | 0.00422 |
| hsa-miR-548n | -3.42309 | 0.00617 |
| hsa-miR-520h | -3.7024 | 0.00059 |
| hsa-miR-570-3p | -4.24066 | 0.00503 |
| hsa-miR-25-3p | -4.2431 | 0.01372 |
| hsa-miR-496 | -4.45449 | 0.00087 |
| hsa-miR-942-5p | -4.58922 | 0.00169 |

MicroRNAs with increased and decreased expression in patients with *RYR1*-RD compared with healthy controls. **<u>Hsa-miR-4454+hsa-miR-7975 were targeted by a single Reporter probe hence data presented is a summed expression of both microRNAs (hsa-miR-4454 and hsa-miR-7975) in the sample.</u>**The final thirteen base pairs at the 3’ end of both miRNAs, hsa-miR-4454 and hsa-miR-7975, are identical.

**Figure 1.**
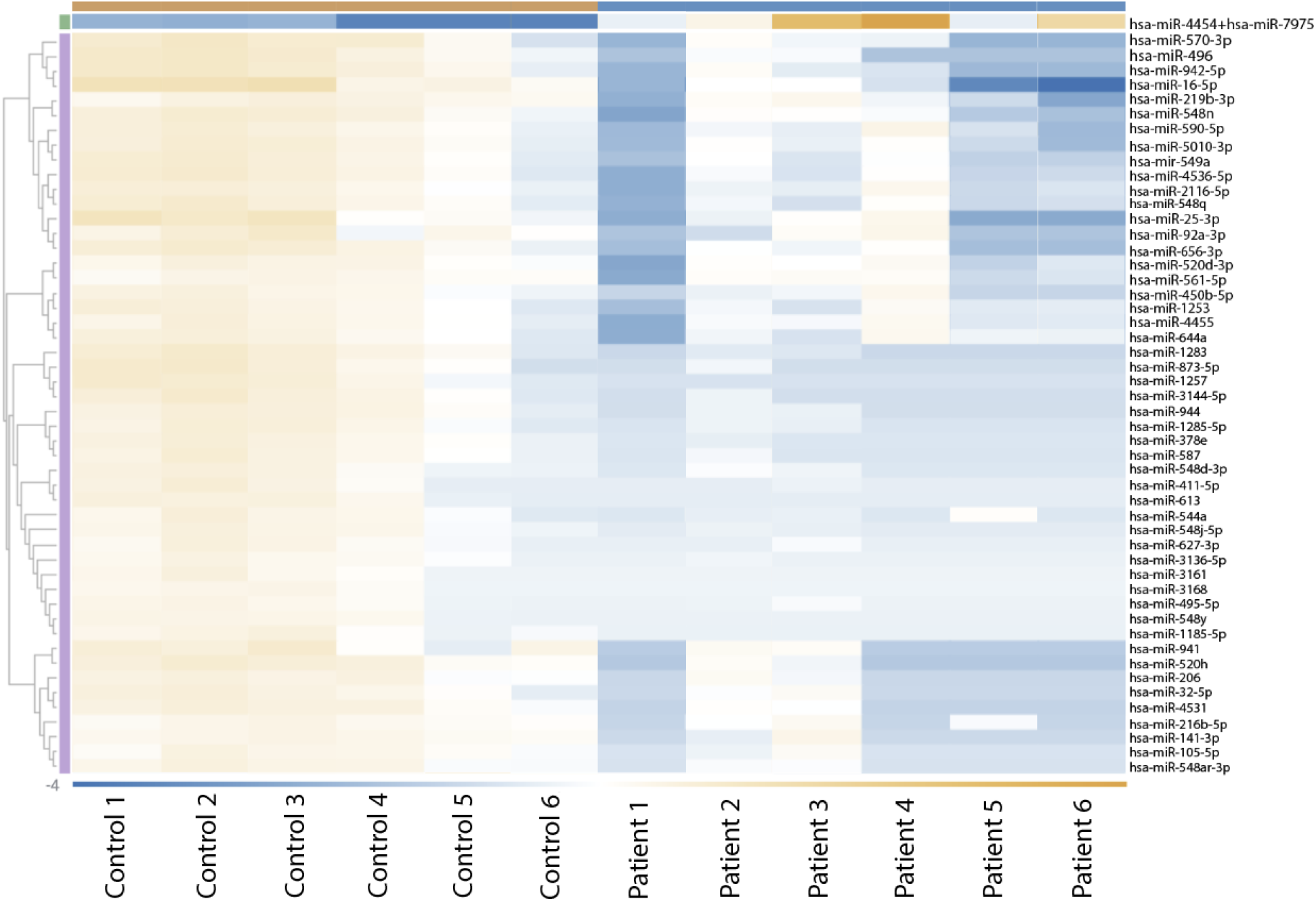
Heatmap of 51 miRNAs in patients with *RYR1*-RD compared with healthy controls. Over-expression is depicted in orange and under-expression is depicted in blue.

**Figure 2.**
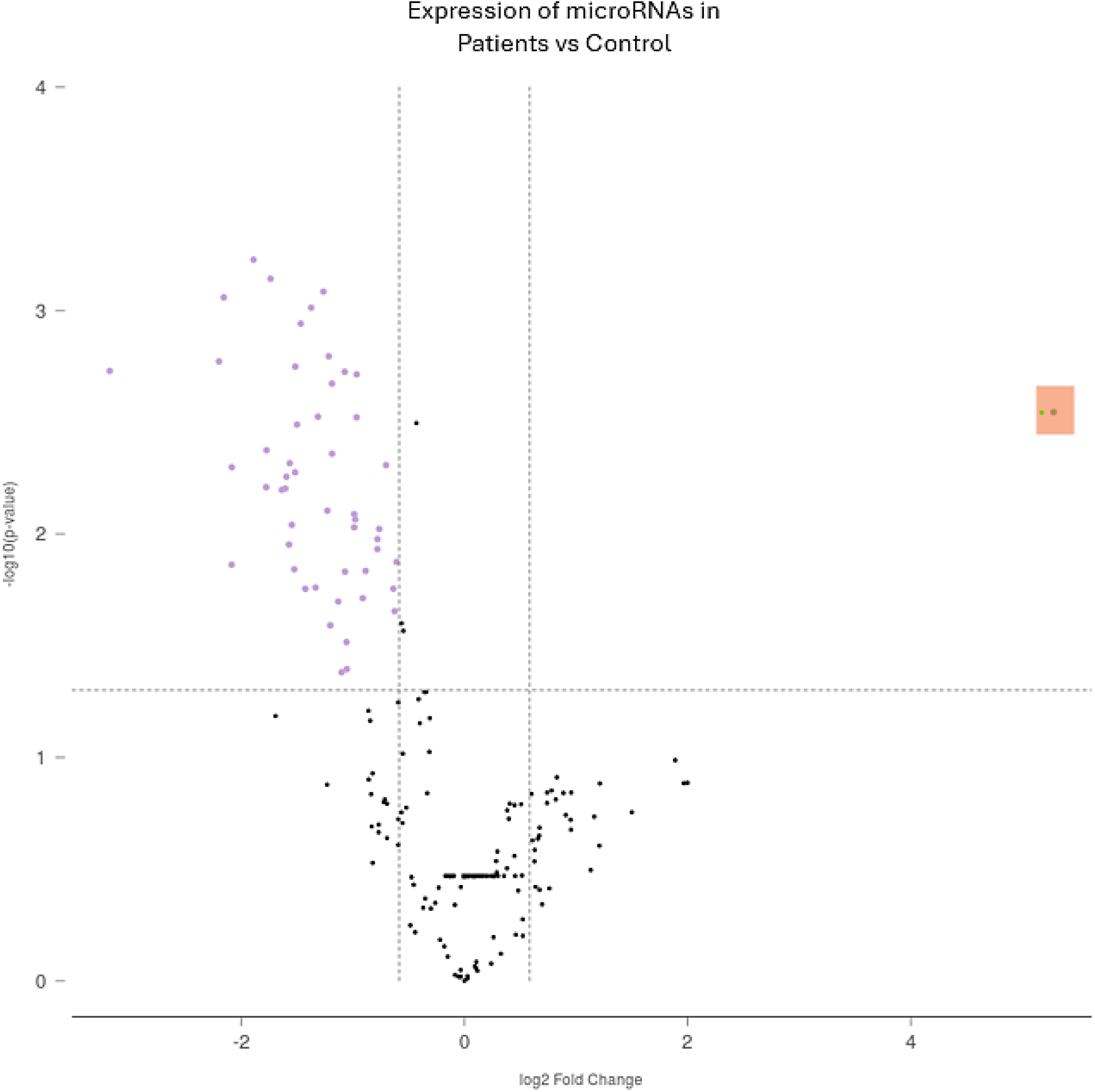
Volcano plot of expression of miRNAs in patients compared with controls. Purple dots represent miRNAs that were under-expressed, green dot (highlighted) represents two miRNAs that were over-expressed (hsa-miR-4454+hsa-miR-7975), and black dots represent miRNAs that were excluded as background levels of expression.

## 4. Discussion

MicroRNAs have emerged as regulators of myogenesis, gene expression, and skeletal muscle function, suggesting that specific miRNAs could be promising clinical biomarkers.^27^ We report on the miR signature of *RYR1*-RD patients using plasma samples from 4 monoallelic and 2 biallelic cases. Of the 51 miRNAs differentially expressed, 49 were under-expressed while two, hsa-mir-4454+hsa-mir-7975 (38.85-fold change, p<0.05), were over-expressed compared to controls.

The final thirteen base pairs at the 3’ end of both miRNAs, hsa-miR-4454 and hsa-miR-7975, are identical. Due to lack of variability, the genes were paired, and a single Reporter probe readout was calculated. The counts from this probe were a summation (+) of the expression of both miRNAs in the sample. It is possible that only one of the two miRNAs is overexpressed in *RYR1*-RD, therefore, further validation is needed. Of note, there is more information on hsa-miR-4454 compared with hsa-miR-7975 in the miRbase database.^28^ However, with the highest fold change of 38.85, the overexpression of hsa-miR-4454+hsa-miR-7975 in *RYR1*-RD presented itself as an interesting target. Both miRNAs have been linked to inflammatory diseases, tumor development, extracellular matrix remodeling, early atherosclerosis (animal study) and reported as a plasma miRNA signature of severe COVID-19.^28 29 30^ In a study assessing the diagnostic value of miRNAs in myalgic encephalomyelitis/chronic fatigue syndrome (ME/CFS), hsa-miR-4454+hsa-miR-7975 was reported to be significantly under-expressed in peripheral blood mononuclear cells and over-expressed in extracellular vesicles.^31^ Although there were no associations found in previous literature between miR-4454+miR-7975 and neuromuscular disorders, predictions of their putative target genes and associated pathways such as calcium signaling, VEGF, and Wnt signaling could be informative. miRNAs can have a number of mRNA targets, it is plausible that *RYR1* is a candidate along with previously reported potential hsa-miR-4454 gene targets such as *CCR4* and *TRAM2*.^32^ Interestingly, *CCR4* is predicted to be involved in Ca^2+^ -mediated signaling. *TRAM2*, a homolog of the translocation-associated membrane protein, is a component of the translocon which is a gated channel through the ER membrane that interacts with the ubiquitously expressed ER Ca^2+^ pump SERCA2b.^33^

The regulatory functions and aberrant expression of miRNAs in neuromuscular disease etiology appear to be disease specific,^34 35 36^ with upregulation correlating with severity of muscle fiber destruction.^7 37^ A previous study investigating the epigenetic effects of muscle-specific (myomiRs) miRs (133, 22, 124, 1) on the regulation of RyR1 protein abundance reported decreased miRs in muscle biopsy samples from biallelic cases with decreased RyR1 protein expression compared with monoallelic cases.^11^ One of the reported myomiR, miR-206, is specifically expressed in skeletal muscle, with higher expression in slow-twitch oxidative fibers,^38^ was not downregulated in their report. Additionally, mixed expression of miR-206 in animal models of Limb-Girdle Muscular Dystrophy (LGMD) have been reported.^7^ However, miR-206 was significantly under-expressed (fold change: -2.48153) in our *RYR1*-RD patient plasma samples consisting of two biallelic cases. miR-206 is essential for cell formation and muscle differentiation during embryogenesis.^39 40^

Up-regulation of miR-206 has been linked to other neuromuscular disorders including Duchenne muscular dystrophy (DMD) and amyotrophic lateral sclerosis (ALS),^41^ with correlations between high miR-206 serum expression levels and low functional performance.^42^ Overexpression of miR-206 inhibits cell cycle progression, while under-expression inhibits cell cycle withdrawal.^43^ As blood miRNAs provide information about the tissues they are secreted from, the data suggest that under-expression of miR-206 in *RYR1*-RD patients is potentially a signature of neuromuscular disorders, though without disease specificity and larger sample validation requirement.

To our knowledge, there are no prior reports on plasma miRNAs in *RYR1*-RD. Our study was limited to a cohort of six *RYR1*-RD patients and six matched controls, thus expanding our patient cohort will be an ongoing focus and further work will be required to characterize the 51 potential signatures, with a focus on hsa-mir-4454+hsa-mir-7975. Additionally, this cohort of patients are from different sub-categories of *RYR1*-RD, a notoriously heterogenous group of disorders. This places reservations on direct comparisons beyond our mono- and bi-allelic sub-classifications. Notable expression patterns observed in our patients suggest that miRNAs emerging from reports such as this one could lead to novel insights and may be used as alternative biomarkers in clinical studies in the future. Our study exhibits how recent advances in multiplex analysis of RNA techniques are expected to accelerate biomarker validation and development in neuromuscular diseases such as *RYR1*-RD. Combined with modalities such as quantitative proteomic analysis of *RYR1* patient samples and metabolomic detection of impaired pathways or elevated metabolites unique to *RYR1*-RD,^44 45^ miRNAs implication as a disease signature provides a dynamic picture of post-translational modifications and an interesting starting point that warrants further exploration and validation.

Development of novel therapeutic strategies and identification of reliable biomarkers that closely correlate with disease severity is crucial for clinical trial readiness. As modulators of cellular phenotypes and skeletal muscle differentiation and proliferation, through sequence-specific interactions with their target mRNA’s 3’ untranslated regions (UTR).^46 47^

## 5. Acknowledgements

This study was funded by the National Institute of Nursing Research (Division of Intramural Research). We thank the RYR-1 Foundation for their continued support and advocacy on behalf of the *RYR1*-RD community. The authors acknowledge Min Mo (Nanostring Technologies) for his support and collaboration during analyses; Jason Sinclair (National Institute of Child Health and Human Development) for his collaboration on this manuscript; NIH Clinical Center 7SWN inpatient unit staff for care of study participants; and NIH Clinical Center Department of Transfusion Medicine for obtaining control plasma samples.

## 5.1 CRediT Statement

Pooja Varma: Validation, Formal Analysis, Investigation, Data curation, Writing-Original Draft/Review & Editing, Visualization. Mayra Saintilus: Resources, Writing-Review & Editing. Morin Nessim: Resources, Writing-Review & Editing. Joshua J. Todd: Conceptualization, Methodology, Resources, Writing – Review & Editing, Supervision, Project Administration. Payam Mohassel: Resources, Writing-Review & Editing. Tokunbor A. Lawal: Conceptualization, Methodology, Resources, Writing – Review & Editing, Supervision, Project Administration, Funding acquisition.

## 7. Disclaimer Statement

This research was supported [in part] by the Intramural Research Program of the National Institutes of Health (NIH). The contributions of the NIH author(s) are considered Works of the United States Government. The findings and conclusions presented in this paper are those of the author(s) and do not necessarily reflect the views of the NIH or the U.S. Department of Health and Human Services.

